# Estimating the energy requirements for long term memory formation

**DOI:** 10.1101/2023.01.16.524203

**Authors:** Maxime Girard, Jiamu Jiang, Mark CW van Rossum

## Abstract

Brains consume metabolic energy to process information, but also to store memories. The energy required for memory formation can be substantial, for instance in fruit flies memory formation leads to a shorter lifespan upon subsequent starvation (Mery and Kawecki, 2005). Here we estimate that the energy required corresponds to about 10mJ/bit and compare this to biophysical estimates as well as energy requirements in computer hardware. We conclude that while the reason behind it is not known, biological memory storage is metabolically expensive,

---

The human brain consumes some 20W of energy, 20% of the body’s total consumption at rest. The cost for computation and information transmission, mostly for synaptic transmission and spike generation, is well documented, and the brain’s design is now widely believed to be constrained by energy needs (Attwell and Laughlin, 2001; Lennie, 2003; Harris et al., 2012; Karbowski, 2012). More recently the metabolic cost of learning has been added to the brain’s energy budget. Experiments in Drosophila indicate that these costs are substantial. In Mery and Kawecki (2005) flies were exposed to a classical conditioning protocol and learned to associate an odor to a mechanical shock. After the protocol, all feeding was stopped and the time to die from starvation was measured. It was found that the conditioning reduced the lifespan compared to control flies. After controlling for exposure to unconditioned and conditioned stimuli separately, the decrease in lifespan was some 20%.

Currently, it is not clear which neural processes are the main energy consumers associated to learning and memory. However, it is known that not all forms of memory are equally costly. Persistent forms, such as Long Term Memory (LTM) in the fly, are costly, but the less persistent Anaesthesia Resistant Memory (ARM) memory which decays in a few days (Margulies et al., 2005), is not. Interestingly, aversive LTM is halted under low energy conditions (Plaçais and Preat, 2013). Such adaptive regulation is also found in mammals where late-phase Long Term Potentiation (late-LTP) is halted under low energy conditions, while early phase LTP is not (Potter et al., 2010).

In this note we review estimates for the energy required to store a few bits of information, namely the association of an odour with a noxious stimulus as happens in the protocol of Mery and Kawecki (2005). The estimates have large uncertainties that will hopefully be narrowed down in the future. Nevertheless, we feel that these ‘ball park’ figures are useful for theoretical considerations and future experiments. We also discuss the estimate in the context of computer hardware.

How much information is stored in the classical odor-shock conditioning? In order to learn the association of odor and shock requires at least one bit of information, namely whether the stimulus is to be avoided or not. If the valence of the stimuli were stored in more detail, a few extra bits would be needed. Furthermore, the animal could store the context of the stimulus, which would be functionally beneficial. However, in contrast to mammals, we have not seen evidence for contextual fear conditioning in flies. We therefore estimate that some 10 bits are stored.

## Direct measurement of energy intake after learning

There are various methods to estimate the energy need for memory formation from experiments. The first method is based on the fact that right after learning, flies increase their sucrose intake to about double the normal rate (Fig1.c in Plaçais et al., 2017). In the Capillary Feeder assay (CAFE), the fly’s energy uptake is determined from the consumption of sugar water from a capillary (5% sucrose; sucrose carries 16.2kJ/g) (Ja et al., 2007). The increase corresponds to an additional intake of 42±160mJ (19±85mJ in Fig 6.e) compared to control flies, where the errors denote standard deviations.^1^

**Figure 1:**
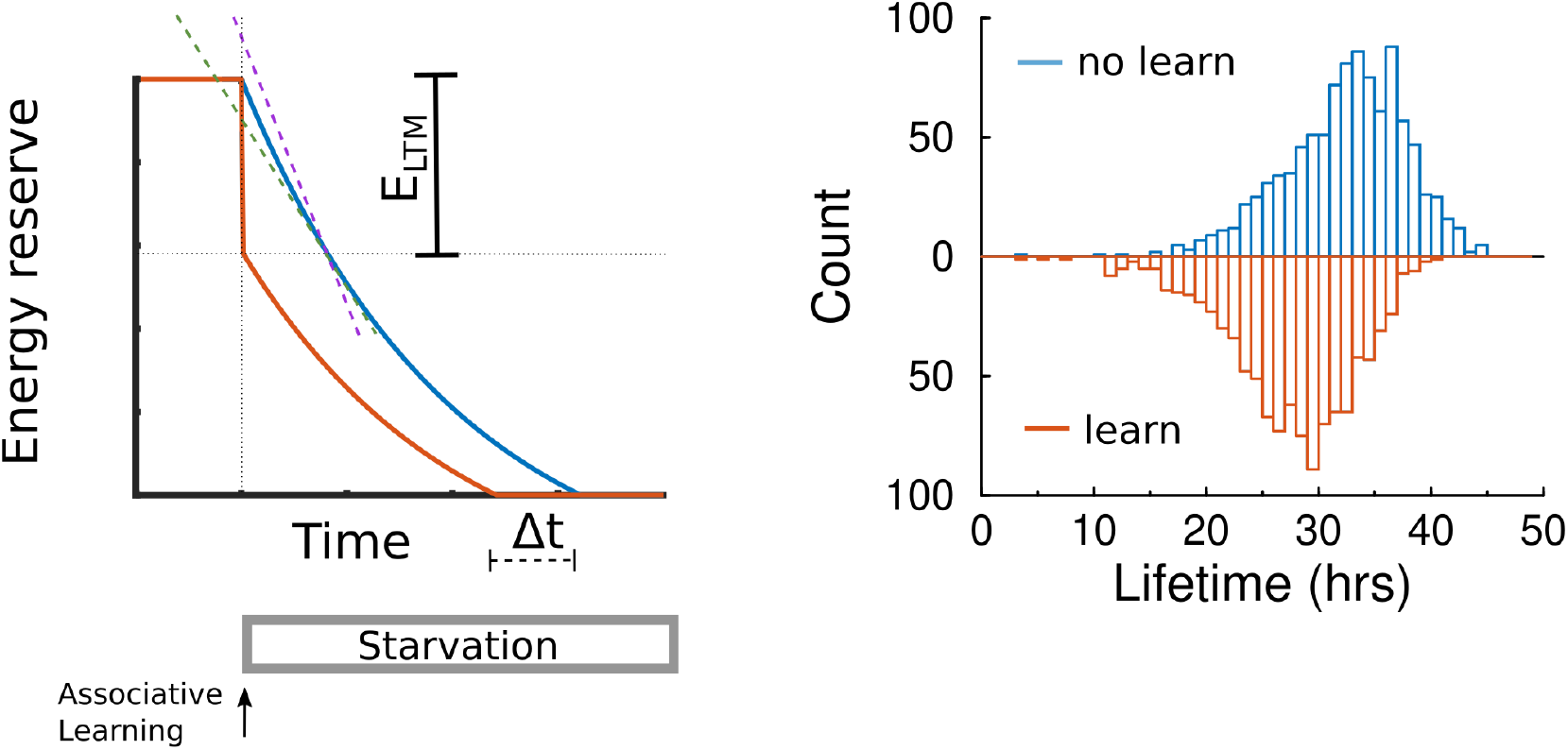
Left: Diagram for estimating the energy used for learning. Energy reserve is plotted over time. At time 0, well fed flies either learn an association, leading to a decrease in energy reserve (orange), or are part of the control group (blue). Subsequently both group are starved and die when their reserve hits zero; taught flies die an interval Δ*t* earlier. The estimated energy used for learning *E*_LTM_ can be estimated from Δ*t* using either the consumption rates right before (purple line) and right after learning (green line). The true value falls between these estimates. Right: Simulation of hazard model for 1000 flies in either the naive (top) or learning population (bottom).

Of this energy intake, some will be lost due to metabolic inefficiency and some will be lost in urine and feces. Assuming a 43% conversion efficiency to produce ATP (Nakrani et al., 2022), one can infer that learning consumed some 20mJ in the form of ATP.

### Estimation via lifetime

An alternative estimate of the energy used for memory formation can be found from the reduction in survival time upon starvation after learning. It is simplest to assume that the fly dies whenever its energy reserve *E*(*t*) drops below zero. Next, assume that the energy reserve decreases linearly in time with a rate *β*. I.e. *E*(*t*) = *E*_0_ – *βt*, where *E*_0_ is the initial energy reserve. Calorimetry can be used to estimate the consumption rate for a non-starving fly at *β* = (7 ±2)*μ*W at 23 C. (Fiorino et al., 2018). (Noting that the metabolic rate varies across strains and that the basal metabolic rate increases steeply with increasing temperature; Klepsatel et al., 2019). The average observed lifetime shortening, denoted Δ*l*, caused by LTM memory formation was about 4.5hrs in both the experiments of Mery and Kawecki (2005) and Plaçais et al. (2017). Thus under this linear decrease model, one finds *E*_LTM_ = *β*Δ*l* = (110 ±100)mJ.

To examine the robustness of this estimate we add realistic features to this model and show how this affects the estimate. First, the energy consumption decreases as reserves are diminishing (Fiorino et al., 2018). That is, the energy reserve is a convex function of time. Fig. 1 left shows the energy reserve versus time in two conditions. In the control condition (blue curve) starvation start at time 0, causing a gradual drop in the reserve. In the learning condition (orange curve) learning causes a rapid drop in the reserve and takes place right before starvation starts.

We assume that metabolic rate is a function of the current energy reserve only. This means that after expending energy on learning, the energy reserve follows the same trajectory as that of a fly that has been starved some time already. In other words, the learning associated expenditure of the energy advances the energy trace by an amount Δ*t*.

We denote the initial rate of consumption as a positive number *β* (slope of purple line; horizontally shifted horizontally for clarity), and that after learning as rate *β*′ (*β*′ ≤ *β*; green line). From Fig. 1, it can be seen that the energy estimate is bounded as

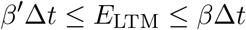

This means that an estimate of the energy cost from lifetime differences Δ*t* based on *β*, is possibly an over-estimate, but that based on *β*′ is an underestimate. The metabolic rate after learning, *β*′, has to our knowledge not been measured directly, however in the setup of Fiorino et al. (2018) the metabolic rate drops some 30% under a calorie restricted diet.

The calculation also holds when the energy consumption caused by learning is not instantaneous as long as *β*′ is measured after the additional consumption caused by learning has stopped.

### Hazard model

A more involved model to estimate the energy consumed by learning is to use a hazard function formulation. A hazard function describes the instantaneous probability for dying at a certain energy reserve level (see e.g. Modarres et al., 1999; Gerstner and Kistler, 2002). In the hazard formulation, even if a population of flies all start with the same energy reserve, they will die at different times. The most basic example is a constant hazard. In that case the lifetimes are exponentially distributed and the mean lifetime is the inverse of the hazard rate.

We denote the hazard at a given energy reserve by *h*(*E*). The hazard increases as the energy reserve drops. We assume that the starvation experiments are so drastic that any age dependence of the hazard can be ignored. (Note that inclusion of age dependence would lead to further underestimation of the energy – in the extreme case that life time is only age dependent, large changes in *E*_LTM_ will not affect lifespan).

In general, the mean lifetime *l* in a hazard model is given by

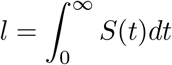

where the survival function *S*(*t*) is given by 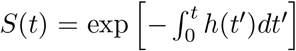. We explore how advancing of the energy trace due to learning as in Fig.1 changes the average lifetime. With a tilde we denote the quantities after learning. The advance means that 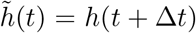, so that the survival function for the learning flies is

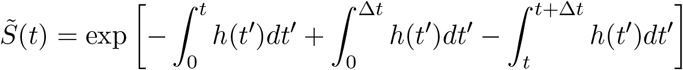

For small Δ*t* this can be approximated as

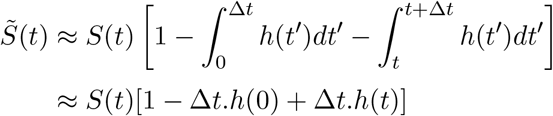

The average lifetime of the learned fly is 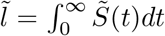. Using that 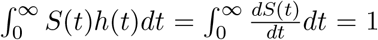, the lifetime after spending an amount *E*_LTM_ at time zero is reduced to

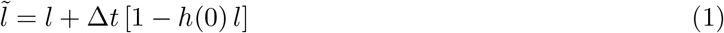

It is instructive to study the two limiting cases: When the hazard is independent of energy and hence constant in time (*h*(*t*) = *h*(0)), one has 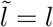. In that case there is no change in lifetime. In the other case, when there is no hazard before starvation, that is, *h*(0) = 0, one has 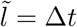. In general the lifetime change 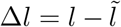 will range between 0 and the shift in the energy profile Δ*t*. Combined with the above result,

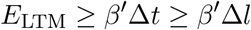

Thus using consumption rate *β*′, one will always *underestimate* the energy expended on learning. The hazard formulation always exaggerates the underestimate. With the caveat of the unknown rate *β*′, we conclude that *E*_LTM_ ≳ 100*mJ*.

As an illustration of this model we simulated 1000 flies, Fig.1b. The initial reserve was set to 0.6J and we assumed it decayed exponentially as *γdE*/*dt* = –*E* – *c*, where *γ* = 40hrs and *c* = 0.3J. The hazard was modeled as *h* = exp(–*kE*)/hr, with *k* = 20*J*^-1^. This resulted in a lifetime of 32.3 hrs without learning, and 27.6hrs when learning. The estimated expenditure (*β*′Δ*l*) was 95mJ, compared to a true value of *E*_LTM_ =100mJ used in the simulation.

## Discussion

In summary, we used two ways of estimating the amount of energy needed to learn a simple association from behavioural data, namely from excess sucrose consumption and from change in lifespan. Encouragingly, the estimates yield comparable numbers on the order of 100mJ, or some 10mJ/bit. It is interesting to compare these costs to memory costs in digital computers. Both data storage and data transmission from CPU to memory cost substantial amounts of energy. In typical personal computers the slowest, most persistent, and most energy costly storage is farthest removed from the processor (Das et al., 2015). For instance, a typical modern Solid State Drive (SSD) can write up to 3GByte/s, taking about 10W (Samsung 970). Hence the energy cost of storage on an SSD is about 0.5 nJ/bit. A hierarchy of smaller and faster caches (L3, L2, L1) speeds up read and write access of data that is repeatedly used by the CPU. Energy costs of these are only of the order of pJ/bit (Molka et al., 2010; Das et al., 2015). With the caveat is that computers are highly optimized for processing large chunks of data, memory storage in computers is therefore some 6…7 orders of magnitude less costly than biological memory storage.

Why is biological learning so metabolically demanding? Currently it is not clear whether most energy is consumed on the synaptic level, network level, or organism level. The biophysical cost of synaptic plasticity in mammals was estimated by Karbowski (2019). The leading cost there is by far protein phosphorylation, which far outweighs estimates for protein synthesis, transport costs and other costs such as actin thread milling. It is estimated as 3 ×10^6^ ATP/synapse/min. Hence the cost of increased phosphorylation in a single synapse during 1 hour would come to 9pJ. Even with 1000 synapses undergoing plasticity this number is still 3 orders of magnitude below the behaviour based estimates above. Moreover, phosphorylation is more characteristic of early, inexpensive early phase LTP than of the expensive late phase LTP. Interestingly, there is recent evidence for different metabolic pathways for different types of plasticity (Dembitskaya et al., 2022). So while those estimates are thus not inconsistent with our estimates, a large amount of energy use remains unaccounted for.

To determine if the missing energy is used directly by synaptic plasticity, it would be of interest to measure energy consumption when the number of modified synapses or the number of memoranda is varied. If learning two associations would costs double the energy, synaptic processes are likely the main consumer. In that case the energy needed to learn multiple associations could rapidly become enormous. For instance learning the well-known MNIST data set requires at some 10^8^ synaptic updates (in preparation). Saving strategies will be needed in that case (Li and Van Rossum, 2020).

An alternative is that the major consumers are changes in brain activity by coordinated processes such as replay – which contributes to memory consolidation in mammals, but also in flies (Cognigni et al., 2018). Calorimetry during learning could provide insight into such hypotheses. Finally, behavioral or physiological changes resulting from the learning protocol might explain the increased consumption. Control experiments with unpaired stimuli in Mery and Kawecki (2005) might not have completely corrected for such effects.

No matter the answer, animals are likely constrained by the high metabolic cost of learning; their savings strategies will help to understand biological memory formation.

## Acknowledgments

We would like to thank Pjotr Dudek and William Levy for discussion. Jiamu Jiang is supported by a Vice-Chancellor International award from the University of Nottingham.

## Footnotes

1 When comparing across experiments it should be noted that in the CAFE assay, energy consumption is strongly reduced during an initial habituation period during the first few days (Van den Bergh, 2022).

